# Gene model correction for PVRIG in single cell and bulk sequencing data enables accurate detection and study of its functional relevance

**DOI:** 10.1101/2022.11.02.514879

**Authors:** Sergey Nemzer, Niv Sabath, Assaf Wool, Zoya Alteber, Hirofumi Ando, Amanda Nickles-Fader, Tian-Li Wang, Ie-Ming Shih, Drew M. Pardoll, Sudipto Ganguly, Yaron Turpaz, Zurit Levine, Roy Z. Granit

## Abstract

Single cell RNA sequencing (scRNA-seq) has gained increased popularity in recent years and has revolutionized the study of cell populations; however, this technology presents several caveats regarding specific gene expression measurement. Here we examine the expression levels of several immune checkpoint genes, which are currently assessed in clinical studies. We find that unlike in most bulk sequencing studies, PVRIG, a novel immune-modulatory receptor in the DNAM-1 axis, suffers from poor detection in 10x Chromium scRNA-seq and other types of assays that utilize the GENCODE transcriptomic reference (gene model). We show that the default GENCODE gene model, typically used in the analysis of such data, is incorrect in the PVRIG genomic region and demonstrate that fixing the gene model recovers genuine PVRIG expression levels. We explore computational strategies for resolving multi-gene mapped reads, such as those implemented in RSEM and STARsolo and find that they provide a partial solution to the problem. Our study provides means to better interrogate the expression of PVRIG in scRNA-seq and emphasizes the importance of optimizing gene models and alignment algorithms to enable accurate gene expression measurement in scRNA-seq and bulk sequencing. The methodology applied here for PVRIG can be applied to other genes with similar issues.

## Introduction

PVRIG is a novel immune-modulatory receptor in the DNAM-1 axis (Alteber, Kotturi, et al., 2021; Whelan et al., 2019a). PVRIG directly inhibits T and NK cell activation while also competing with the co-activating receptor DNAM-1, for the binding of a shared ligand, PVRL2, which is expressed on tumor and antigen-presenting cells. An antibody blocking the PVRIG/PVRL2 interaction was shown to increase CD8+ T-cell and NK cell activity in preclinical studies (Whelan et al., 2019a), and is currently being evaluated in early clinical trials [NCT04570839, NCT03667716]. The advancement of scRNA-seq methods has enabled a more accurate characterization of PVRIG expression and of other drug targets within different cell types. Consequently, we have shown that PVRIG expression is uniquely enriched on early-differentiated T stem-cell memory (TSCM) CD8+ T cells (Alteber, Cojocaru, et al., 2021), unlike other immune checkpoints such as CTLA4, PD1, LAG3, TIM3, and TIGIT which are more highly expressed on exhausted CD8+ T cells (Wherry & Kurachi, 2015). This unique expression pattern in TSCM suggests a potential for improved clinical response, as these cells have been shown to possess an increased capacity to expand and self-renew to generate waves of activated T-cells (Klebanoff et al., 2005; Krishna et al., 2020), and their increased levels were found to be associated with a positive response to immunotherapy treatment (Sade-Feldman et al., 2018).

In recent years droplet-based single-cell RNA sequencing has gained increased popularity within the scientific community, mainly utilizing the 10x Genomics (10x) Chromium platform (Svensson et al., 2020). In support of this platform 10x have developed the CellRanger software suite which allows easy and uniform conversion of raw sequencing data into a cell-by-gene count matrix (Zheng et al., 2017). One of the innovations implemented in droplet technology is the use of 3’ or 5’ biased mRNA sequencing which allows considerable cost saving (less than $1 per cell) and thus enables the profiling of tens or hundreds of thousands of cells; greatly promoting the study of cell populations and tissue composition (Zhang et al., 2019). However, probing the expression of individual genes using this technology might pose challenges associated with the relative sparsity of the data (Wang et al., 2021).

## Results

### PVRIG is poorly detected in 10x chromium scRNA-seq data

While studying the expression of T-cell checkpoints in scRNA-seq data, unexpectedly, we found that PVRIG expression in droplet-based data (most commonly 10x Chromium) is often considerably lower versus what is observed in full-length scRNA-seq (e.g., well-based Smart-Seq) or in bulk RNA-seq (Figure 1a,b) and does not align with flow cytometry data for surface protein expression (Whelan et al., 2019a). We hypothesized that the observed reduced expression of PVRIG in existing scRNA-seq datasets is a technical artifact stemming from droplet-based scRNA-seq. The literature describes three potential culprits for reduced gene expression in droplet-based scRNA-seq: (1) reads mapping immediately 3’ to known gene boundaries due to poor 3’ UTR annotation; (2) intronic reads stemming from unannotated exons or pre-mRNA; (3) discarded reads by read count software (e.g. CellRanger, RSEM) due to multi-gene mapping to paralogous genes or read-through transcripts. We thus examined mapped reads within and around the PVRIG gene locus using bulk RNA-seq, 3’ and 5’ droplet-based scRNA-seq, and long-read scRNA-seq from PBMC and sorted T cells data (Figure 1c,d). We did not observe a substantial number of reads downstream of the 3’ end of the gene, suggesting that the cause is not an unidentified longer 3’ UTR (Figure 1d). We also did not find an unusually high number of reads mapped to the intronic regions of the gene or to other loci in the genome. In contrast, we identified an overlapping transcript, STAG3-209 (ENST00000451963) that overlaps the entire PVRIG gene region (Figure 1c,d). This transcript is curated under the GENCODE gene model, which relies on Ensembl transcripts, and is the default used by CellRanger. STAG3-209 stems from a readthrough of the STAG3 gene upstream of PVRIG, its supporting evidence consists of only one sequence originating from testis tissue (Figure S1a,b), and it is predicted to undergo non-sense mediated decay (NMD) (ENST00000451963.1 Ensembl Entry). Importantly, the weak supporting evidence of STAG3-209 resulted in the exclusion of this transcript from the RefSeq gene model (Figure S1c).

**Figure 1:**
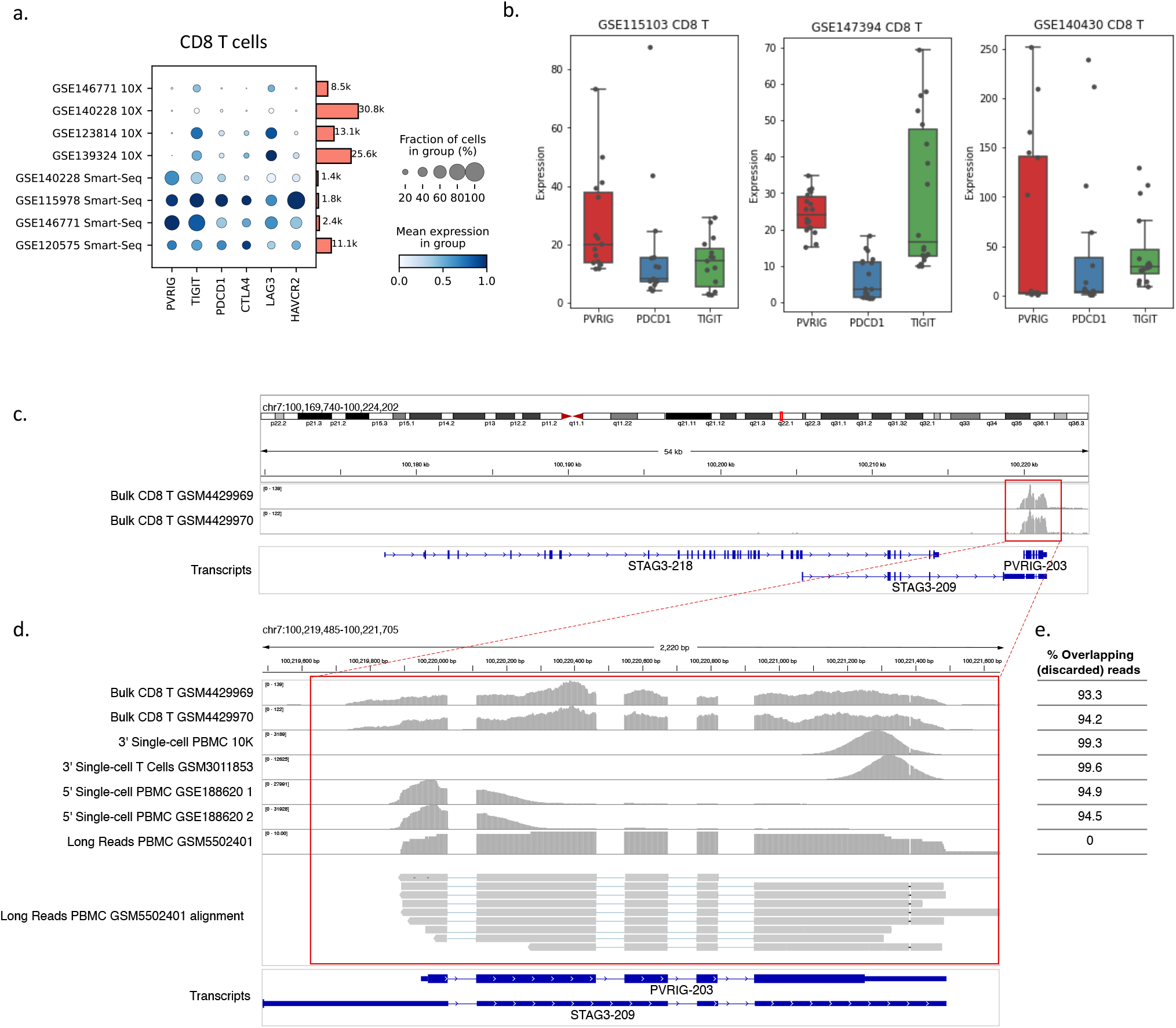
PVRIG is poorly detected in 10x Chromium data and overlaps with STAG3-209. **a**. Dotplot showing the expression of PVRIG and additional T-cell checkpoints in several 10x datasets (top) versus Smart-seq2 datasets (bottom) collected from TME. Bars indicate number of cells in each sample. **b**. Boxplots showing the expression levels of PVRIG, TIGIT and PD-1 across several CD8 T-cell datasets as measured using bulk RNA-seq. GSE140430 isolated CD8 T-cells from TME, rest are from blood. **c**. The STAG3-PVRIG genomic locus. Histograms feature the distribution of reads in bulk RNA-seq data of isolated CD8 T-cells highlighting that PVRIG is expressed in such cells (red box). Gene structures appear below in blue. **d**. Closer view of PVRIG locus. Histograms highlight the distribution of reads in the indicated datasets. **e**. Fraction of reads that map to both PVRIG and STAG3-209 and are thus discarded.

### Little evidence supports STAG3-209, suggesting it could be discarded

The exons and introns of the canonical PVRIG (PVRIG-203) are almost entirely identical to that of the overlapping predicted STAG3-209 except for the second PVRIG intron, 82 bp length, and a slightly shorter (by two bp) 3’ end of STAG3-209 (Figure 1d). We thus examined the mapping of reads from several bulk RNA-seq of sorted CD8 T cells around the second intron of PVRIG. We found that in all samples the reads follow the exact gene structure of PVRIG and almost no reads (except stochastic noise-level reads) are mapped to the intron that is part of the predicted STAG3-209 transcript, whereas several reads in each sample are mapped to the junction between exon 2 and exon 3 in PVRIG (Figure 1c). Thus, at least in CD8 T cells, all reads mapped to the PVRIG-STAG3 overlap are most likely to originate from the PVRIG mRNA. In addition, we examined exon-exon junctions from the cancer genome atlas (TCGA), data spanning over 12K samples, and could not detect junctions that support the expression of the STAG3-209-PVRIG readthrough while junctions supporting the PVRIG gene were readily found (data not shown).

Because most short reads do not fall within exon-intron junctions, we also examined long-read data based on PacBio sequencing of T-cells (Tian et al., 2022). We found that all reads spanning the second intron of PVRIG do not contain the retained-intron which is found in STAG3-209 and there are no junctions that support STAG3-209 (Figure 1d). Thus, at least in these datasets, there is no support for the expression of STAG3-209.

### Gene model correction enables accurate detection of PVRIG expression

In light of different technical issue that can potentially hinder the expression of specific genes in scRNAseq, Pool et al recently proposed a modified gene reference model in which readthrough transcripts were removed; including deletion of STAG3-209 (Allan-Hermann Pool et al., 2022). In order to verify the specificity of the change we also generated a gene model in which we removed only the STAG3-209 readthrough and validated that only PVRIG expression is altered by the modification (Figure S2). To test the effect of the modified references, we ran the standard CellRanger pipeline using these improved gene models on 10x Chromium public dataset released by 10x which contains PBMCs as well isolated CD8 T-cells from 4 donors that were confirmed by bulk RNA-seq to express PVRIG (Figure 2, Figure 1b). Indeed, when we used the improved gene models, we found elevated PVRIG expression in CD8^+^ T cells and NK populations previously found to express it (Figure 2; Whelan et al., 2019). As expected, expression of other T cell checkpoints was not altered by the correction as expected. Moreover, the differential expression of PVRIG across cell types following the correction reflects protein levels of PVRIG previously assessed by flow-cytometry in human normal and tumor tissue (Figure 2) (Whelan et al., 2019a). To further validate our correction, we generated scRNA-seq profiles of immune populations isolated from the TME of patients with ovarian cancer. These samples had been previously analyzed by flow cytometry as PVRIG-positive (Figure 3). We found that following the correction PVRIG demonstrated solid expression levels and patterns that match those previously described, namely high expression in NK and TSCM cells (Figure 3 a,b).

**Figure 2:**
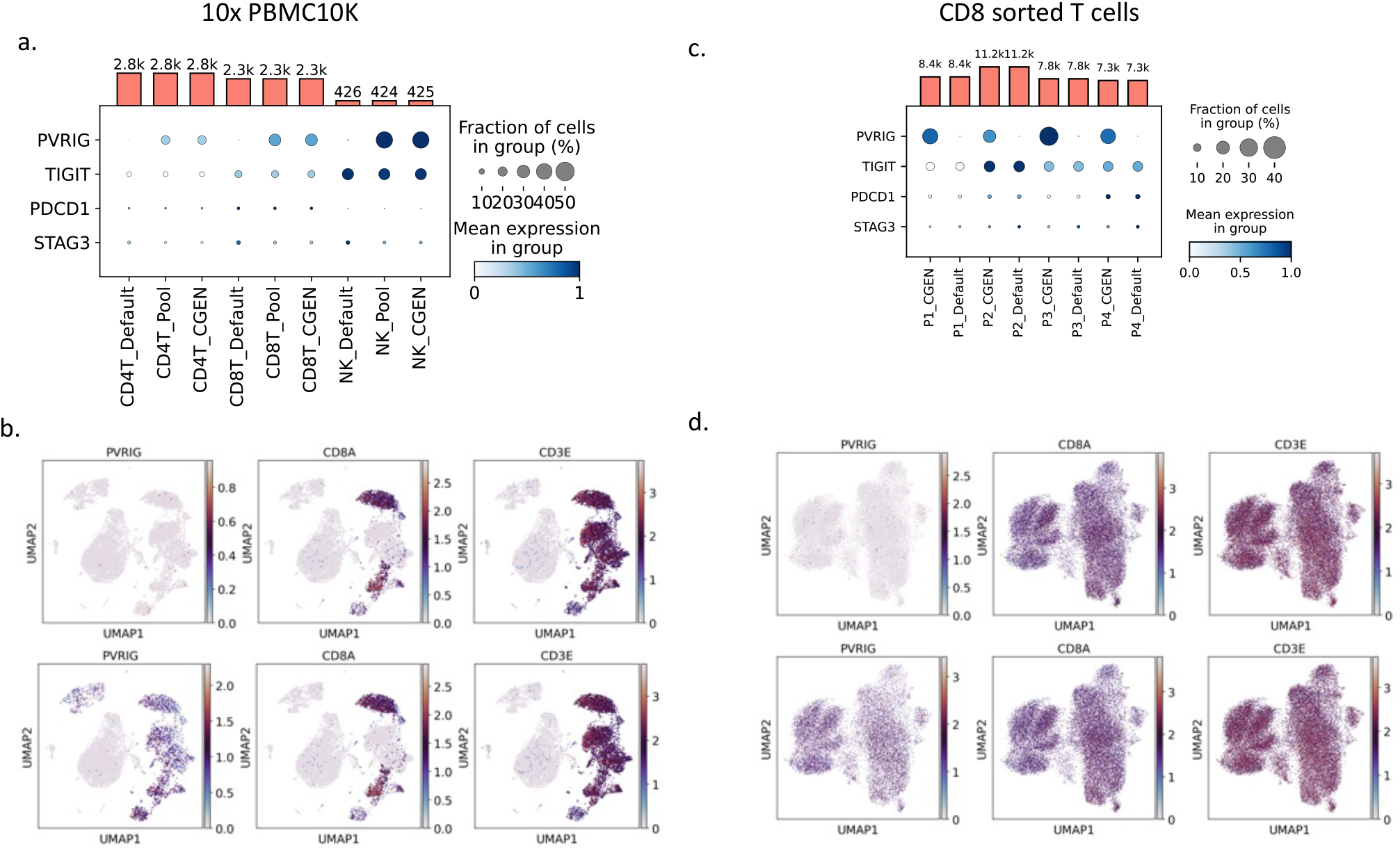
Gene model correction allows proper detection of PVRIG. **a**. Dotplot showing the expression of PVRIG and additional T-cell checkpoints as well as STAG3 in 10x PBMC10K dataset processed using various gene models subset for T and NK cells. Default – GENCODE, Pool – previously published, broadly corrected reference. CGEN – gene model sepecifically corrected in the PVRIG gene locus. Bars indicate number of cells in each sample. **b**. UMAP showing the expression of PVRIG, CD8A and CD3E in the 10x PBMC10K dataset using the default gene model (top row) and the corrected gene model (bottom row). **c**,**d** – Same as a,b but for CD8+CD95+ sorted T-cells from 4 healthy donors (GSE147397).

**Figure 3:**
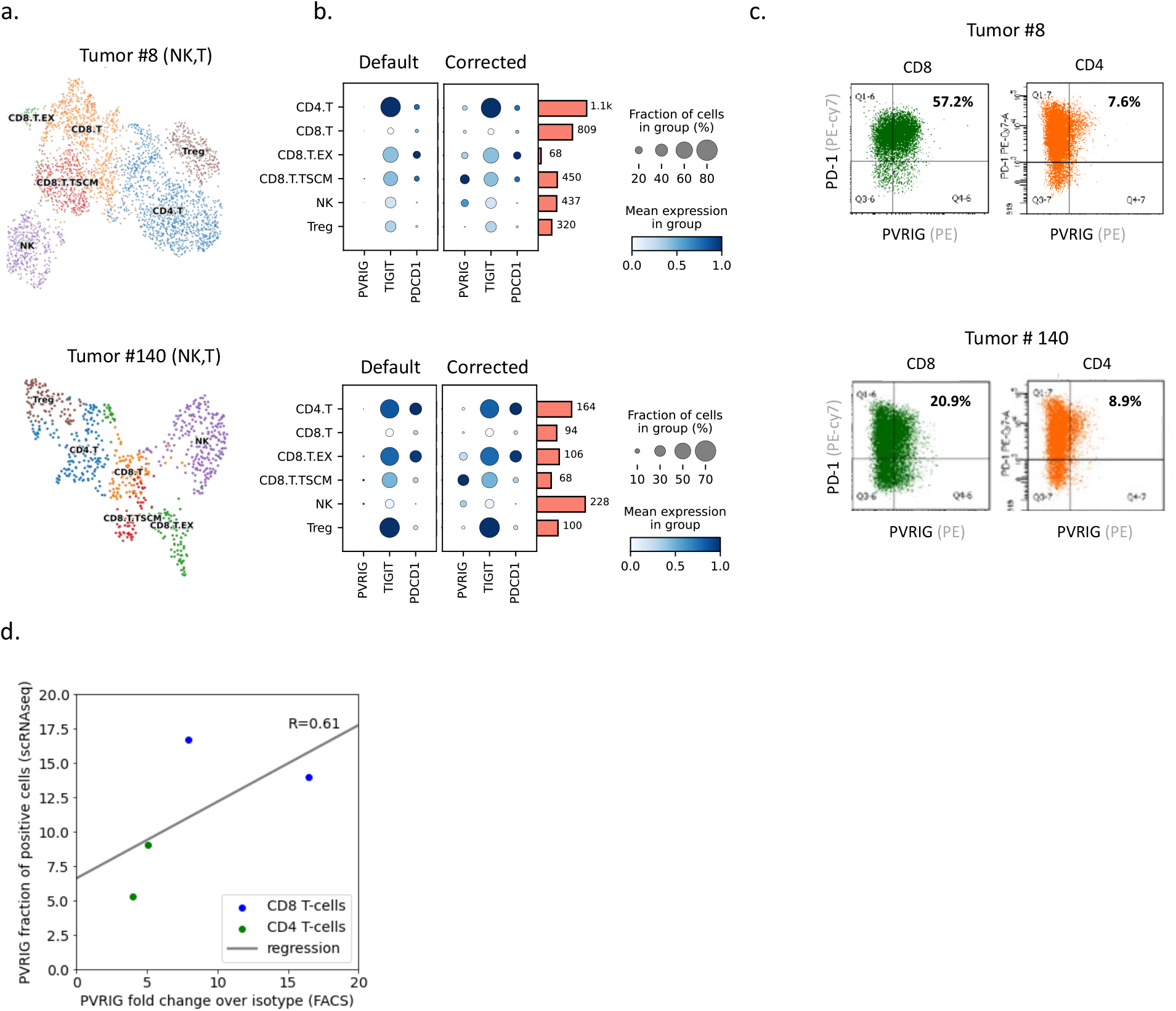
Corrected PVRIG expression in ovarian tumor samples demonstrates a correlation with protein expression. **a**. UMAPs showing the population distribution in two ovarian tumor samples subset for T and NK cells. **b**. Dotplot showing the expression of PVRIG and additional T-cell checkpoints in ovarian tumor samples processed using the default or corrected gene model subset for T and NK cells. Bars indicate number of cells in each sample. **c**. Matched FACS analysis showing the expression of PVRIG and PD-1 in CD8 or CD4 T cells. **d**. Correlation between PVRIG-positive fraction as measured using FACS (X-axis) to its parallel in the scRNA-seq data following the gene model correction. Linear-regression line is shown in gray as well as the Pearson correlation coefficient.

Furthermore, we noted that the mRNA levels of PVRIG correlated with the protein levels (though with low number of supporting samples) measured by FACS in cells obtained from the same patient-derived samples (Figure 3c,d). Together, our findings suggest that the discarding of reads by CellRanger due to multi gene mapping was responsible for the observed decreased expression.

### Gene model correctness determines accurate expression in bulk RNA-seq data

Considering our findings, we were interested to uncover the underlying cause which allows PVRIG to be readily detected in Smart-seq2 and bulk sequencing while it is poorly detected in 10x Chromium data. Hence, we analyzed bulk sequencing using the STAR aligner, which is used by CellRanger, and using the GENCODE reference or our corrected gene model and found that PVRIG could only be detected upon using our model (Figure 4a). Thus, we concluded that the underlying reason for poor detection of PVRIG does not stem from 3’ or 5’ bias used in 10x droplet-based single cells data, but due to the gene model used by default which contains the STAG3-209 transcript. We have examined several publications that demonstrated PVRIG expression in Smart-seq2 and noted that they indeed employed gene models that do not contain STAG3-209.

**Figure 4:**
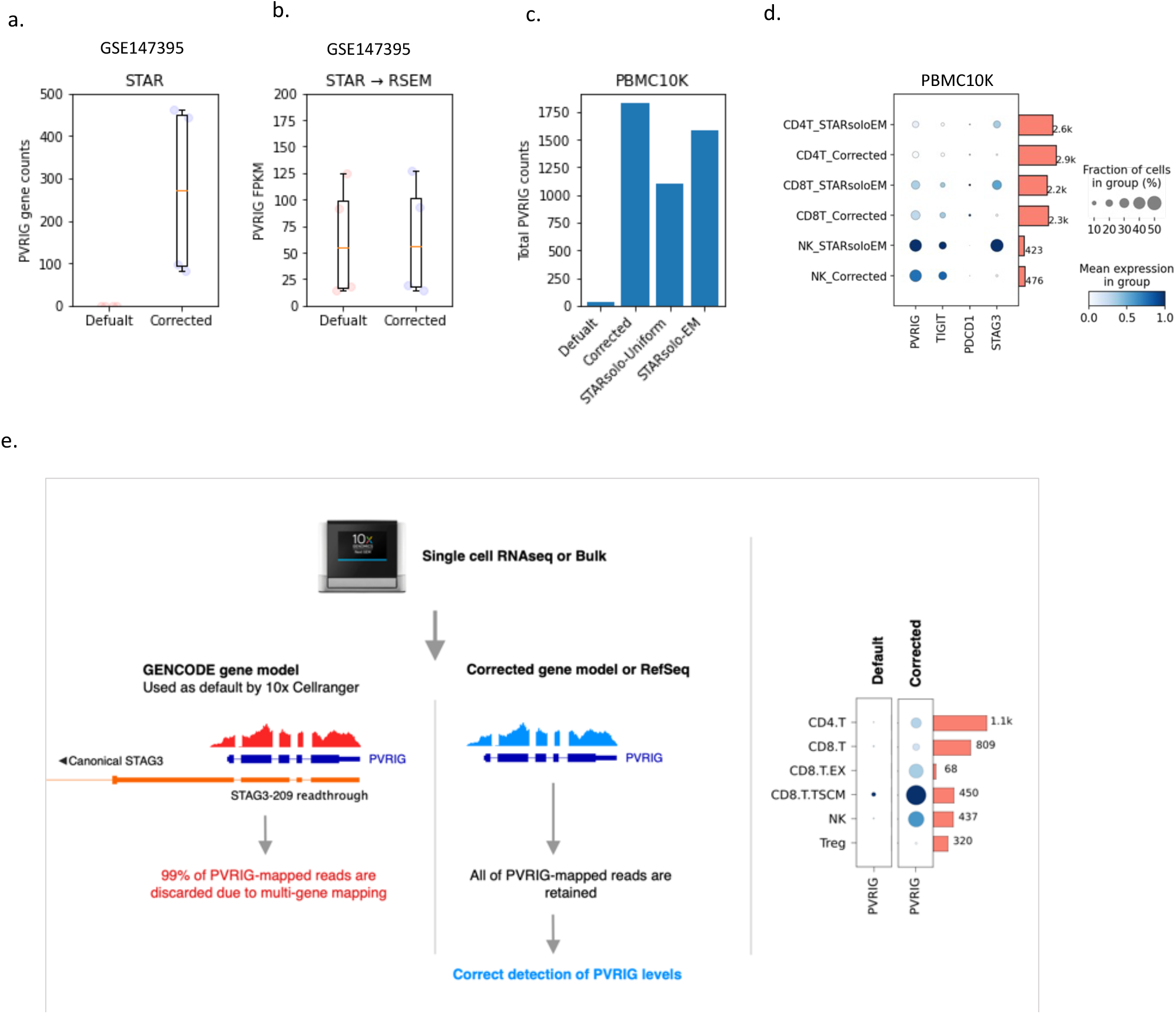
Correction using multimappers and summary illustration. **a**. Boxplot showing the number of PVRIG read counts per sample in bulk sequencing data of CD8 T-cells using the default or corrected gene model. **b**. Same as in a, but following multimap correction using RSEM. **c**. Barplot showing the number of PVRIG counts in PBMC10K dataset processed using the default, corrected gene model or following multi-mapping correction using STARsolo. **d**. DotPlot showing the expression of PVRIG, TIGIT, PD-1 and STAG3 in NK/T populations in the PBMC10K dataset processed using the corrected gene model or using STARsolo-EM method. **e**. Summarizing illustration; using the GENCODE gene model reads are discarded due to multi-mapping, following the correction of the gene model PVRIG expression is correctly measured.

### Multimap algorithms partially rescue the expression of PVRIG

The issue of multi-mapped reads was extensively studied for full-length RNA-seq data, and algorithms that distribute the multi-mapped reads between their alignments based on the uniquely mapped read ratio have been developed (Deschamps-Francoeur et al., 2020). We therefore analyzed bulk data of CD8 T-cells using the RSEM read estimator (Li & Dewey, 2011), that employs STAR aligner data as input and contains algorithms to handle multi-mapped transcripts. We noted that indeed RSEM was able to recover PVRIG gene expression in bulk RNAseq (Figure 4b) regardless of the gene model used. More recently, efforts were made to resolve the issue of multi-mapped reads in scRNA-seq data as well (Benjamin Kaminow et al., 2021; Srivastava et al., 2019). To test the feasibility of such algorithms to correct the PVRIG-STAG3-209 multi-mapping we ran STARsolo on data which was aligned to the default gene model, using expectation maximization (EM) or uniform distribution as means to correct multi-gene mapping. We noted that the tool was indeed able to improve the detection of PVRIG, to similar levels obtained by employing gene model correction (Figure 4c,d). Further expression analysis demonstrated that the multi-map correction applied using STARsolo also increased the expression of STAG-3 in a way that is likely to be an artifact (Figure 4d). Though STARsolo was able to improve the expression of PVRIG it is apparent that this feature is not yet fully supported as the package currently generates a non-filtered matrix which holds the multi-mapping data, forcing the user to conduct additional non-trivial processing steps.

## Discussion

One of the strengths of scientific investigation is the great diversity of approaches and highly distributed manner by which results are repeated using different methodologies. 10x Chromium revolutionized the field of scRNA-seq, allowing a uniform analysis of the outputs, which greatly contributes to the reproducibility of the results across different laboratories. However, as we demonstrate in our study, the decisions made by these tools, such as the selection of reference gene models, could have broad implications. We show that gene models should not be taken for granted when studying the expression of specific genes and emphasize the importance of refining such models. Our work reveals that PVRIG expression is underestimated in many single-cell, and potentially also in bulk, datasets due to technical reasons that stem from the usage of the GENCODE gene model; that includes the rare STAG3-209 readthrough which almost entirely overlaps with PVRIG. While we point the spotlight on PVRIG, due to our specific scientific interest, similar phenomena are likely to affect additional genes as well. As the field of scRNA continues to evolve it is essential for the scientific community to address the inclusion of rare readthrough transcripts and correct annotation of 3’ UTRs. While a holistic, genome-wide, solutions such as the one suggested by Pool et al may be useful, they still require further verification and refinement as they have broad implications on the expression of many genes.

Alternately, algorithms for handling multi-mapped reads count could be integrated into CellRanger, yet these also require rigorous testing and, as we show in our study, currently available tools require further development before they become standard. Meanwhile, we suggest employing our generated gene model which we demonstrated to faithfully correct the expression of PVRIG in a specific manner that does not alter the expression of additional genes. Using our corrected reference, we show that PVRIG expression levels in CD8 T-cells located at the periphery and in primary tumor samples are comparable to other immune-checkpoints, such as TIGIT and PD-1. Thus, we believe that studies which profiled T-cells using 10x Chromium or other methods that have used the GENCODE reference should re-evaluate the expression of PVRIG and other genes that might suffer from a similar phenomenon. Future works and cell atlases should also carefully consider which gene model is utilized. This will hopefully lead to further establishment and understanding of the role of PVRIG in the context of immune-oncology and will support ongoing clinical efforts.

## Methods

### scRNA-seq processing

Single-cell RNA-seq raw data was downloaded from GEO and processed using CellRanger 7.0.0 count option employing three gene model references: 1) GENCODE (default), 2) Model from Pool et al., and 3) a specific model where only STAG3-209 transcript was removed (CGEN model). For Smart-seq2 raw data was not available and thus processed count matrices were used. CellRanger output data or count-metrics were explored and further processed using scanpy 1.9.1 while employing standard preprocessing steps and clustering. Doublets were removed using Scrublet 0.2.3 with default parameters. Clusters were classified based on the expression of known markers (CD8 T-cells: CD3E^+^CD3D^+^TRAC^+^CD8A^+^CD4^-^, CD4 T-cells: CD3E^+^CD3D^+^TRAC^+^CD4^+^CD8A-, Treg: CD3E^+^CD3D^+^TRAC^+^CD4^+^CD8A^-^FOXP3^+^, NK: CD3E^-^CD3D^-^NRC1^+^). For multi-mapping analysis (STARsolo) STAR 2.7.10a was used employing either Uniform or EM options; to generate filtered cells matrices ‘STAR -- runMode soloCellFiltering’ was used. Long-read sequencing (PacBio) was aligned using STAR.

### Bulk RNA-Seq processing

Bulk RNAseq data was downloaded from GEO and processed through STAR 2.7.9a and RSEM 1.3.3 (STAR-based) using the two gene model references as described above.

### Generation of corrected gene model

To generate a corrected gene model the reference provided as default by 10x, GRCh38 (GENCODE v32/Ensembl 98), was downloaded. Rows relating to STAG3-209 were deleted from the GFT file and new reference was generated using ‘cellranger mkref’ for CellRanger scRNAseq or using ‘STAR --runMode genomeGenerate’ for bulk and STARsolo analysis.

### Flow cytometry and scRNAseq

Tumor biospecimens from patients with high-grade serous ovarian cancer (HGSOC) were processed within 6 hours of surgical resection. Dissociation was done using human Tumor Dissociation Kit (Cat # 130-095-929, Miltenyi Biotec, Bergisch Gladbach, Germany) and gentleMACS™ Octo Dissociator with Heaters as per manufacturer’s instruction. PBMCs were enriched by density-gradient centrifugation with Ficoll-Paque Plus (17-1440-02, Cytiva).

Flow cytometry for PVRIG and TIGIT expression on viable CD8 and CD4 T cells was done as previously described (Whelan et al., 2019b). For sorting prior to scRNAseq, 10^6^ cells per sample were stained with the Live/Dead aqua viability dye (Invitrogen Cat# L34957), followed by Fc receptor blocking. Cells were then stained with FITC-conjugated anti-human CD45 (clone HI30, BioLegend). Viable CD45+ cells were sorted on a BD FACSAriaTM Fusion sorter.

Approximately 16,500 immune cells per patient were loaded on a 10x Genomics Chromium platform for targeted recovery of ∼10,000 cells. Libraries were sequenced on a NovaSeq 6000 system.

## Data Availability

The public scRNAseq datasets used in the study: GSE146771, GSE140228, GSE123814, GSE136324, GSE140228, GSE115978, GSE120575, GSE188620, GSM5502401 and 10x PBMC10K (*10k Human PBMCs, 3’ v3*.*1, Chromium Controller*). Bulk RNAseq datasets: GSE115103, GSE140430, GSE147394, GSM4429969, GSM4429970.

Corrected gene model is available for download at https://doi.org/10.5281/zenodo.7274284

## Acknowledgements

We thank Eran Ophir for reviewing the manuscript and insightful comments.

## Supplementary figures

**Figure S1:**
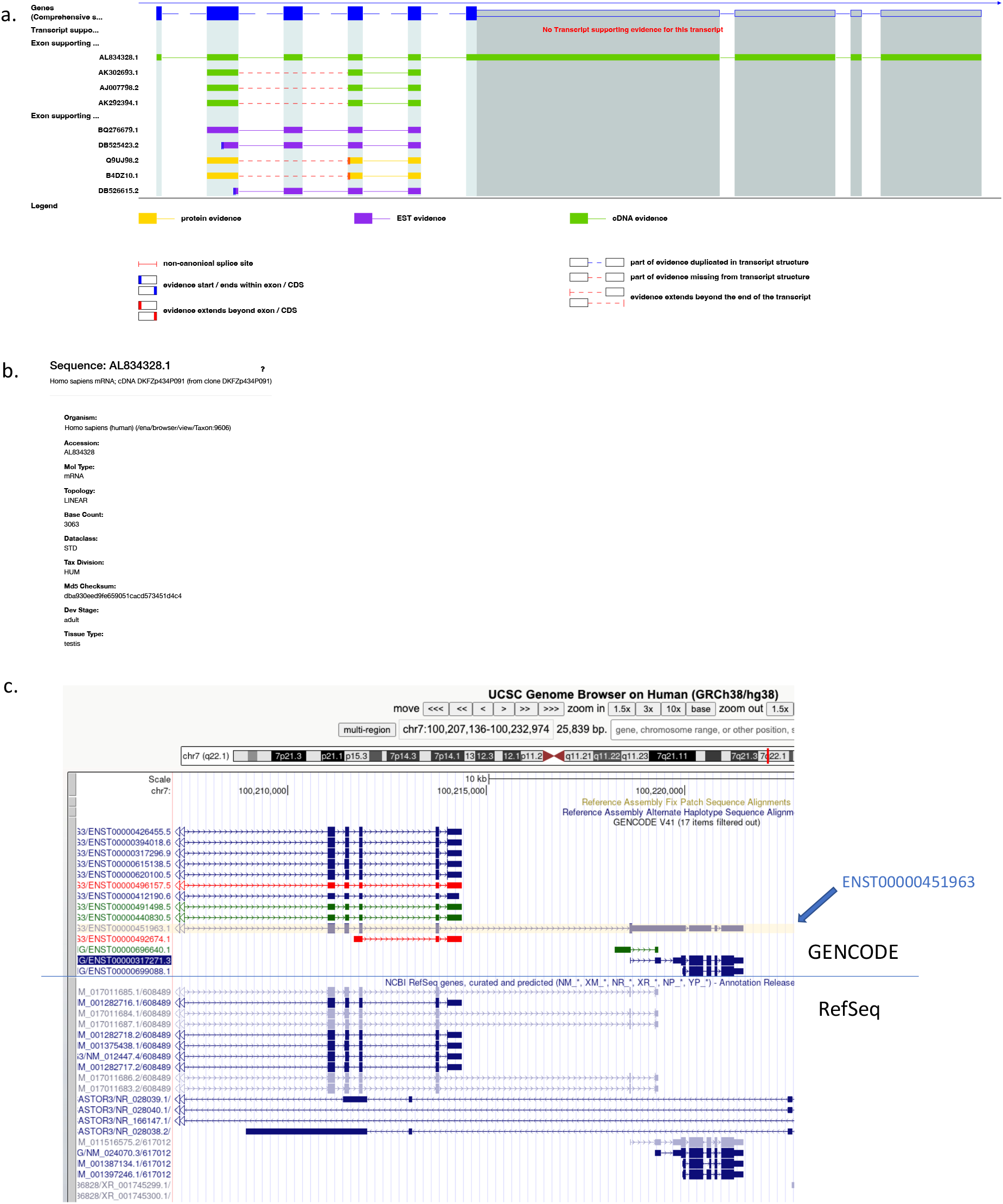
STAG3-209 is a rare transctipt curated by GENCODE and not by RefSeq. **a**. The only supporting evidence for STAG3-209, cDNA with the accession number AL834326.1. **b**. details of the cDNA accession in a. **c**. UCSC genome browser screenshot showing that STAG3-203 is curated by GENCODE and not by RefSeq.

**Figure S2:**
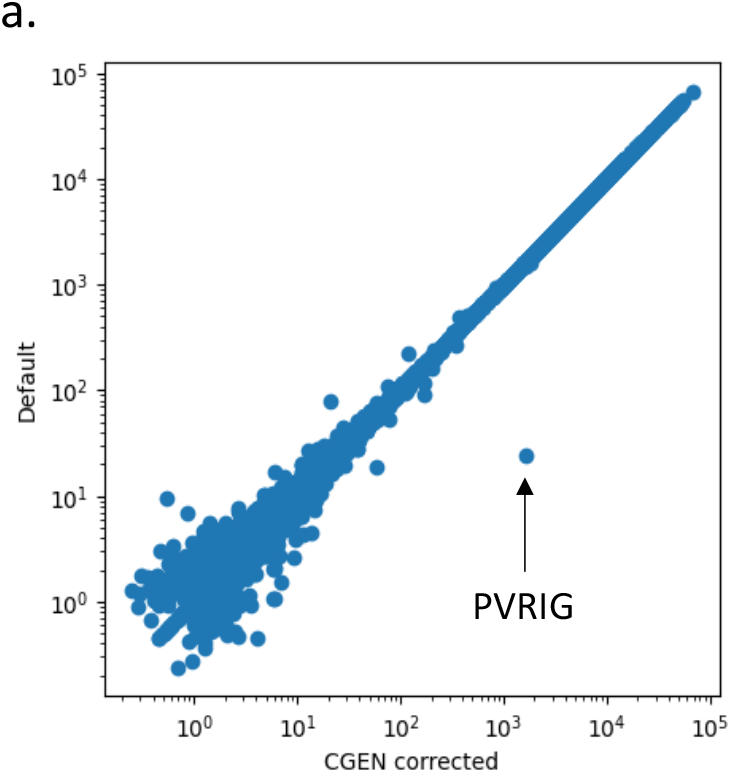
PVRIG specific gene model correction vsersus genomewide correaction. **a**. Scatterplot showing the read counts for individual genes in the default gene model and following the deletion of STAG3-209 (CGEN corrected). Arrow highlighting the change in expression of PVRIG.

